# Converting genetic information to non-equilibrium cellular thermodynamics in enzyme-catalyzed reactions

**DOI:** 10.1101/2023.09.15.557926

**Authors:** Robert A. Gatenby

## Abstract

Living systems use genomic information to maintain a stable highly ordered state far from thermodynamic equilibrium but the specific mechanisms and general principles governing the interface of genetics and thermodynamics has not been extensively investigated. Genetic information is quantified in unitless bits termed “Shannon entropy”, which does not directly relate to thermodynamic entropy or energy. Thus, it is unclear how the Shannon entropy of genetic information is converted into thermodynamic work necessary to maintain the non-equilibrium state of living systems. Here we investigate the interface of genetic information and cellular thermodynamics in enzymatic acceleration of a chemical reaction S+E→ES→E+P, where S and E are substrate and enzyme, ES is the enzyme substrate complex and P product. The rate of any intracellular chemical reaction is determined by probability functions at macroscopic (Boltzmann distribution of the reactant kinetic energies governed by temperature) or microscopic (overlap of reactant quantum wave functions) scales - described, respectively, by the Arrhenius and Knudsen equations. That is, the reaction rate, in the absence of a catalyst, is governed by temperature which determines the kinetic energy of the interacting molecules. Genetic information can act upon a when the encoded string of amino acids folds into a 3-deminsional structure that permits a lock/key spatial matching with the reactants. By optimally superposing the reactants’ wave functions, the information in the enzyme increases the reaction rate by up to15 orders of magnitude under isothermal conditions. In turn, the accelerated reaction rate alters the intracellular thermodynamics environment as the products are at lower Gibbs free energy which permits thermodynamic work (*W*_*max*_ *= −ΔG*). Mathematically and biologically, the critical event that allows genetic information to produce thermodynamic work is the folding of the amino acid string specified by the gene into a 3-dimensional shape determined by its lowest energy state. Biologically, this allows the amino acid string to bind substrate and place them in an optimal spatial orientation. These key-lock are mathematically characterized by Kullback-Leibler Divergence and the interactions with the reaction channel now represent Fisher Information (the second derivative Kullback-Leibler divergence), which can take on the units of the process to which it is applied. Interestingly, Shannon is typically derived by “coarse graining” Shannon information. Thus, living system, by acting at a quantum level, “fine grain” Shannon information

## Introduction

Living systems, maintain a stable ordered state while far from thermodynamic equilibrium through acquisition, storage, and translation of information (1-4). McIntosh (5) expressed this simply in a word equation:

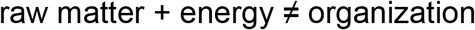

but

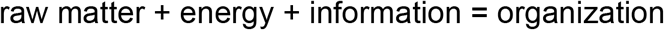

However, the fundamental principles and specific mechanisms governing translation of heritable (genetic) information into a low-entropy, non-equilibrium thermodynamic state (2) remain largely unknown. Here we examine the interface of biological information and thermodynamics through the actions of enzymes: S + E→ ES → E + P, where S stands for substrate, E for enzyme, ES is enzyme substrate complex and P product.

Here we propose general principles and specific mechanisms by which genomic information is translated into a stable low entropy, non-equilibrium state that is a unique property of living systems. Multiple studies have quantified the information content in the genome using Shannon “entropy” (17-19). However, converting Shannon entropy to thermodynamic entropy in living systems is far more challenging than it first appears. The fundamental problem illustrated by conservation of units of measurement. Shannon “entropy is derived from probability function so that metric of information (bits) is unitless (i.e., contains no physical quantities) so that there is no clear theoretical or mathematical mechanism for translation to thermodynamic work.

We approach this problem by focusing on the broader topic of information theory. In 1922, Robert Fisher (6) pioneered the development of information theory. Fisher focused on the quantity of information that is acquired (and, as well, the amount of information lost) in the process of a measurement. This approach is broadly used in virtually every scientific discipline including biology(7), psychology(6), physics (8) and chemistry(9) in a measurement Unlike Shannnon information, because Fisher Information is derived from physical measurement, it assumes the units of the associated physical processes (20-24).

Critical to our analysis here, Shannon and Fisher Information are related (8) as the former emerges from course graining (transitioning from a continuous to a discrete function dx →Δx) of the latter. Thus, because genetic information is encoded as discrete quantities (i.e. the triplet nucleotide code that specifies different amino acids) it is appropriately characterized by Shannon information. Thus, living systems must convert Shannon information encoded in the genome to Fisher Information that can be applied to thermodynamics through a process of fined graining (25-27). To examine this process, we investigate the sequence of steps in which information encoded in a gene forms an enzyme that alters the physical interaction of reactants to produce a chemical reaction.

Normal function in living systems requires acceleration of critical chemical reactions (10). In non-living systems, reaction rates (in the absence of a catalyst) are governed by temperature and an activation energy barrier inherent to each reaction. As defined by the Arrhenius equation, increasing system temperature accelerates reactions by increasing the kinetic energy of the reactants and, thus, the probability they will overcome the activation energy. However, living systems are typically constrained to narrow temperature ranges limiting their ability to use “thermodynamic” acceleration of reaction rates through an influx of heat. Instead, living systems employ heritable (genomic) information to generate enzymes that accelerate chemical reactions by up to 15 orders of magnitude (11).

While enzymes have a degree of flexibility that may allow them to impact the reactants with increased kinetic energy, current conceptual models of enzyme function emphasize their role in capturing the reactants and placing them in close proximity (12) and precise orientation to resemble the reaction transition state (11) with minimum separation of their quantum wave functions (von Neumann entropy (13)). This was described by Pauling as an “enhanced transition state (14)” and may include quantum tunneling (15).

Thus, in living systems, chemical reactions may be accelerated by two mechanisms (Figure 1): 1. An influx of heat that increases the temperature and, therefore, the kinetic energy of the reactants as described by the Arrhenius equation and 2. Optimization of the proximity and spatial orientation of reactants by catalysts, which allow quantum interactions (16) that increase the probability of a reaction. The latter was mathematically described by Knudsen over 100 years ago and supported by theoretical and experimental (17-19) evidence.

**Figure 1.**
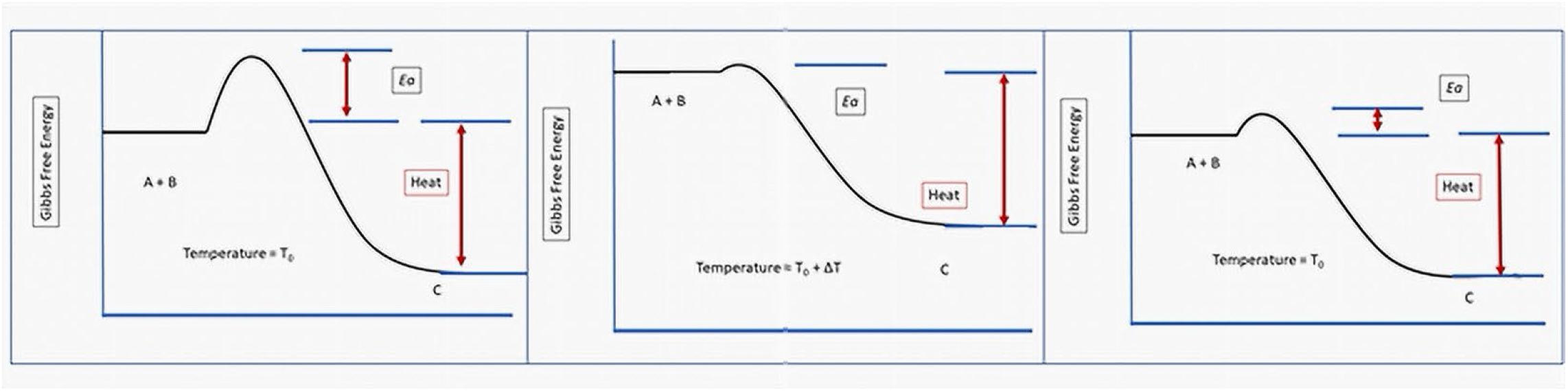
The dynamics of accelerating the chemical reaction A + B → C through injection of heat. The y axis is the Gibbs free energy and the x axis is “reaction coordinates”. Left panel, shows conventional diagram showing the reaction can spontaneously occur because the Gibbs free energy of the product is lower than that of the reactants. The reaction rate is governed by the activation energy, which is a fixed constant that is independent of the thermodynamic state of the system. Middle panel, the reaction is accelerated by the influx of heat which raises the energy of both the reactants and product. Because the activation energy is not changed by the increased temperature, the reaction is accelerated. However, the energy states of the reactants and products are higher. Right panel, addition of an enzyme has the effect of reducing the activation energy but, since an influx of heat is not required, the reaction is accelerated without any change in energy levels of the reactants or product. Thus, the final state of both systems that accelerate a reaction may have the same concentration of reactants and products, its thermodynamic state is quite different.

Here we investigate the parallel thermodynamic (temperature) and biological (enzyme) pathways that control the reaction rates. This equivalence results in a reaction rate expression: exp (−*E*/⟨*E*⟩)cos(*θ*/< *θ*>), that includes thermodynamic (energy) *E* and substate quantum interaction angle *θ*from which both the Arrhenius and Knudsen laws are derived. Thus, a reaction rate is maximized when *either* (a) the reactant kinetic energy *or* (b) the quantum orientation *θ*is optimized.

Although these mechanisms seem quite different, we note both processes can be described by probability functions (Figures 2 and 3). That is, the energy of reactant molecules at any given temperature is governed by a Boltzmann distribution. The probability an impact between reactants will exceed the activation energy (*E*_*a*_) is dependent on these distributions (Figure 2). Similarly, the probability of a reaction is also determined by the quantum orientation (*θ*) of the reactants at the point of collision (Figure 3). Note that these dynamics occur at an interface between molecular (thermodynamic) and atomic (quantum) scales. Importantly, such mixed state systems can be derived by the principle of Maximum Fisher information (20, 21).

**Figure 2.**
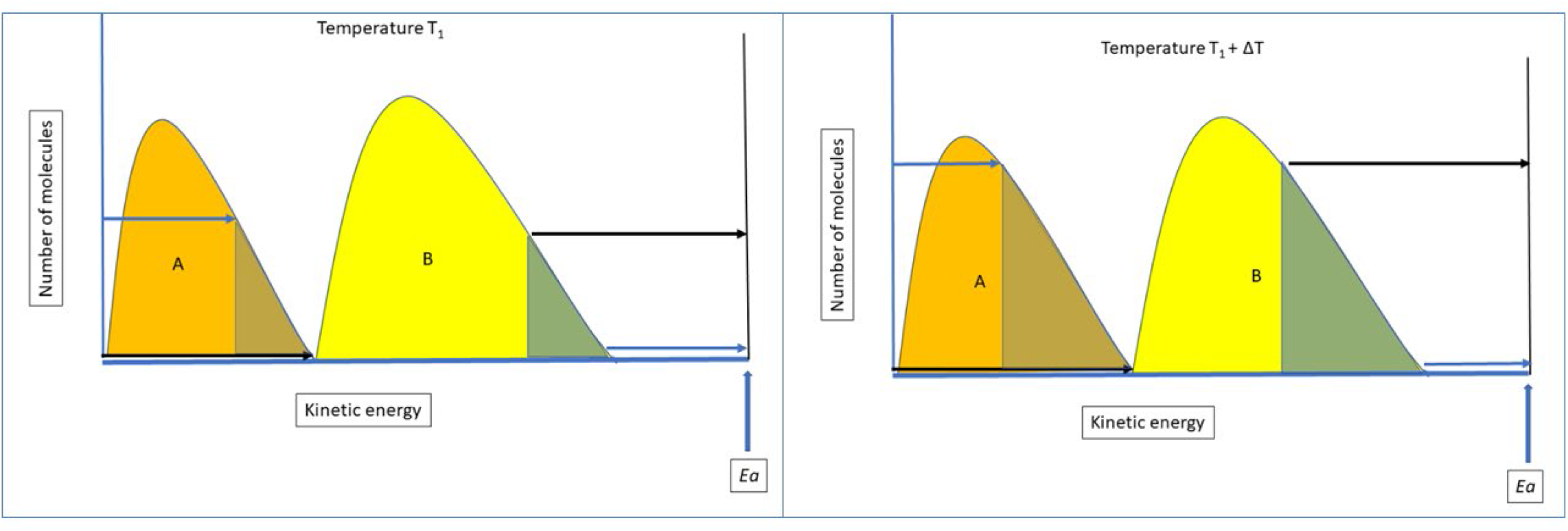
A spontaneous chemical reaction is possible when the Gibbs Free energy of the products is lower than that of the reactants. The rate of reaction increases with temperature, which is interpreted as a threshold minimum kinetic energy (activation energy) of the reactants necessary for the reaction to proceed. (Left panel) A simplistic representation of the activation energy as an additive function of the Boltzmann distribution of the substrate molecules. The activation energy threshold is possible only for a small fraction of the substrate molecules (triangles) depending on the kinetic energy of the other substrate molecule. That is, the additive kinetic energy of two interacting can exceed *Ea* only through interactions of molecules in the two triangles. (Right panel) An increase in temperature moves the Boltzmann distribution of both substrate to the right so that probability that an interaction will achieve the activation energy is increased.

**Figure 3.**
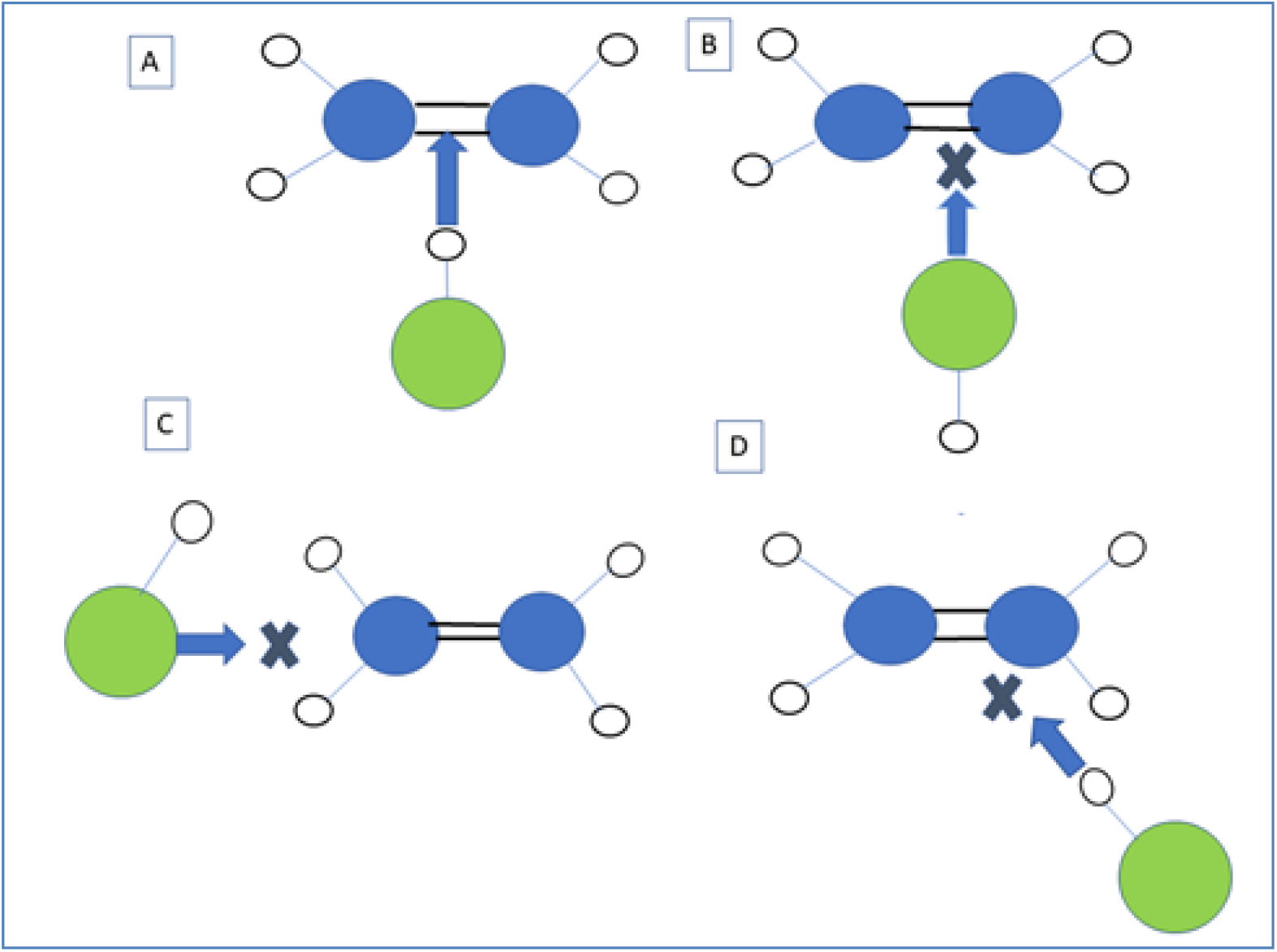
The probability of spontaneous reaction can be viewed as a probability that reactants interact at an optimal orientation to allow quantum interactions at the reaction site. Here, examples of four angles *θ*of orientation between the Ethylene (C_2_H_4_) and HCl molecules. Each case shows a different angle of impact and the corresponding changes in interactions of the asymmetric charge of the HCl molecule and the double carbon bond in ethylene. We arbitrarily assign the vertical axis as the angle that results in a reaction (A). The other angles of interaction (B, C, and D) will not produce a reaction. Thus, the maximum reaction probability is θ = 0, the joint probability law, *p*(***x***) = *p*(*E, θ*), follows.

Thus, we propose Shannon information stored in the genome is translated into a 1-dimensional protein with a specific amino acid sequence. The protein then folds into a 3-dimensional structure that, based on the amino acid sequence, represents a low free energy state. This change in dimensionality and loss of free energy (see Discussion) represents a fine graining of Shannon information. This allows the enzyme to affect the interactions of reactant at a quantum level and an information theoretic transition from unitless Shannon information to Fisher Information which takes on the physical units of the reaction. Furthermore, because Fisher information is conserved in physical processes, including chemical reactions (22, 23), the principle of Maximum Fisher Information (or, equivalently, minimum loss of Fisher Information (24)) it can maximally accelerate the reaction rate without addition of heat. This produces an intracellular state in which the concentrations of substrate and products from several thousand reactions produced by information in enzymes is markedly different from its environment where the reactions are governed by temperature. The rapid reaction rate decreases the concentration of reactants and increases that of the products producing low entropy state as the heat from the reactions are dissipated into the environment. This permits a stable low entropy state that is also far from thermodynamic equilibrium with the environment – properties found uniquely in living systems.

## Results

### Quantum interactions by ‘induced fit’

Let reactants A and B be located within an enzyme at respective spatial coordinates ***x*** and ***r***, with

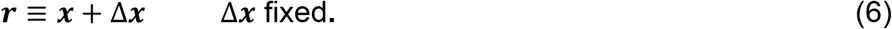

We assume that, during reaction, the reactant B, at *displaced* positions ***r*** = ***x*** + Δ***x***, are spatially and temporally entangled with the reactant A at positions ***x***. Thus, in Eq. (5), p(***x*** + Δ***x***) is entangled with p(***x***). The physical effect is that reactant A positions ***x*** interact, almost simultaneously, with many possible reactant B positions ***x*** + Δ***x***, *all during a single reaction time interval*. A thermodynamic equilibrium is approached whereby -- in analogy with *quantum* realities -- the two reacting molecules A, B are, effectively, two manifestations, or states, of a single product molecule. Hence, we solve problem (5) in this scenario.

### Molecular interactions satisfying constraints

Each chemical reaction generally has its own constraint terms leading to a predicted probabilistic contribution *p*(*E, θ*). Minimizing the loss of information in Eq. (5) promotes the chemical reaction by (a) by lowering the required activation energy of the reaction and (b) bringing together the reactants A and B at *optimal orientation angles θ*promoting bonding of molecules A and B.

The solution obeying principle (1), (5) of maximum Fisher information, found in Appendix A, is

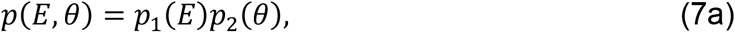

where

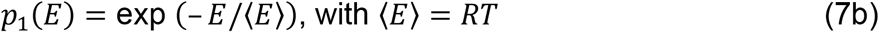

and

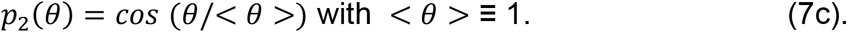

Eqs. (7a-c) describe the Arrhenius-Knudsen “combined” reaction distribution *p*(*E, θ*). Constant parameter *R* is either the gas- or Boltzmann constant, depending on the reaction medium. By Eq. (7a) the reaction phenomenon *p*(***x***) ≡ *p*(*E, θ*) is *the product of an exponential law on* energy *E with a cos(θ)* law on interaction angle *θ*, the Knudsen law, respectively, with

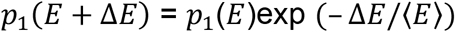

and

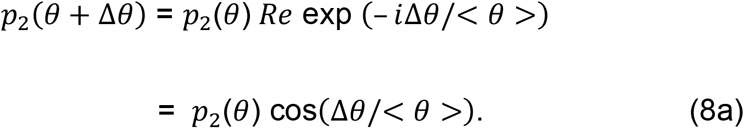

For convenience, normal *and* trigonometric variables are often treated *together* covariantly as a joint *statistical* effect, with a complex exponential on angle replaced, as here, by the *real* cosine of that angle.

From this viewpoint, our result Eqs. (7a-c) of joint Knudsen scattering for variables (*E, θ*) is a *generalization* from a single exponential (Arrhenius) dependence on the energy *E* to a *joint dependence* on energy *E* and scattering angle *θ*. The cos(Δ*θ*/< *θ*>) dependency in Eq. (8a) also accounts for the angular dependence discussed in Fig. 2.

### Cellular thermodynamic considerations

Prior work has demonstrated that Fisher Information provides a critical connection between non-equilibrium thermodynamics and quantum mechanics (24-27). Here, acceleration of a reaction rapidly consumes limited supplies of reactants and increases the concentration of products, thus reducing the number of local microstates (28). Thermodynamically, this results in increased heat production. However, since there is no barrier to heat flow from the cell to its environment, the net effect is decreased entropy while the cell remains isothermal with its environment.

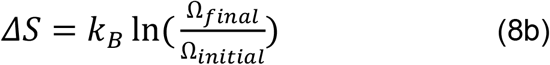

Where *k*_*B*_ is Boltzmann’s constant and Ω is the number of microstates before and after enzymatic acceleration of the reaction. If *Ω*_*initial*_> *Ω*_*final*_ entropy (*S*) decreases; and equivalently, level of order increases.

Thus, the traditional link between thermodynamics and Boltzmann mechanics emphasizes a “top down” dynamic in which a change in thermodynamic state of the system (e.g., an influx of heat causing increased temperature) alters the Boltzmann distribution of the molecular states. We propose biological information alters the Boltzmann distribution by its actions at quantum spatial scales. Thus, the Boltzmann distribution diverges from that expected for the thermodynamic state of the system.

## Discussion

Living systems are stable low entropy states far from thermodynamic equilibrium – a state that is unique in nature. It is intuitively clear that biological information is central to generating this characteristic physical state but the general principles and precise mechanisms by which heritable biologic information alters the thermodynamic state of a cell remains unclear. While genes encode multiple proteins with diverse functions, here we focus on their dynamics as enzymes. We analyze the role of information in intracellular chemical reactions but note other cellular functions such as signaling pathways typically involve enzymatic dynamics such as, for example, phosphorylation and epigenetic changes require methylation and acetylation reactions. Thus, our analysis is also applicable to these diverse cellular functions.

Enzymes allow living systems to accelerate critical reactions without the addition of heat. In a prior study, we demonstrated the genetic information in an enzyme is expressed thermodynamically as a reduction in the activation energy of a reaction (29). Here we investigate the specific mechanism by which enzymes produce a stable, low entropy non-equilibrium state that is uniquely characteristic of living systems.

We demonstrate biological information in the form of an enzyme acts along the spatial continuum of quantum dynamics, Boltzmann statistical mechanics, and thermodynamics. Here, biological information encoded in a molecular sequence and characterized by Shannon information folds into a 3-dimensional structure that represents a local free energy minimum. This process represents a fine graining of the genetic information and allows the information to continuously alter the physical interactions of the substrate. Through evolutionary selection over time, the shape of the enzyme (which is determined by the amino acid sequence) is optimized so that it maximizes the quantum overlap of the substrate allows the reaction to be accelerated without injection of heat. By large acceleration of the reaction rate (up to 15 orders of magnitude), the collective actions of multiple enzymes produce an intracellular Boltzmann distribution with reduced concentrations of reactant and increased products that (because of the reduce Gibbs free energy of the reaction) is lower in entropy while the heat of the reactions (and associated entropy) is dissipated into the environment. Thus, the intracellular Boltzmann state is one that would ordinarily be found at a much higher temperature and, therefore, inconsistent with the thermodynamic state of the environment. That is, biological information produces a stable low entropy state that is far from equilibrium with its environment – the state of living systems that is unique in nature.

The physical connection of thermodynamics and Boltzmann statistical mechanics is well established but most frequently viewed as a “macroscopic → microscopic” process in which thermodynamic perturbations, such as an influx of heat alters the Boltzmann distribution of molecular kinetic energy. In contrast, genetic information functions as a microscopic → macroscopic process. By optimizing quantum interactions between the reactants, biological information accelerates reaction rates and, thus, alters the Boltzmann distribution of the reactants and products. Because the accelerated reaction rapidly converts reactants to products, it reduces the number of available microstates and, therefore, generates a lower entropy. Furthermore, because the environmental temperature is unchanged in the process, heat generated by the reactions dissipates into the environment.

Of particular interest, we note that the fine graining of biological information is the result of a change in dimensionality (folding of a single amino acid string into a 3-dimensional shape) and corresponding loss of free energy. This allows living systems to function at the interface of Newtonian and quantum dynamics. This may represent a fundamental principle of information dynamics and will be the topic of future investigation.

In summary, living systems use microscopic → macroscopic approach, similar to Maxwell’s demon (30), to generate a lower entropy internal state without violating the laws of thermodynamics. Through the additive information of thousands of enzymes, a cell can maintain a highly improbable state that is ordered and stable even while far from equilibrium with its environment. More broadly, our results suggest living systems, through the actions of biological information, function at the interface of the probabilistic quantum and deterministic Newtonian dynamics.

## Methods

### How can information dynamics in chemical reactions be quantified?

The human genome contains about 30 billion bits of Shannon information (31) indicating a high level of order but this unitless metric does not directly translate into chemical or physical processes in living systems.

In contrast, Fisher information (FI) takes on the units of the physical process to which it is applied; and so, has been extensively used in deriving the diverse laws of physics, chemistry, and biology (27, 32-34). Examples include amplitude and phase distributions of quantum mechanics (25), electromagnetics, tumor growth dynamics, covid-19 spread, and distribution laws of thermodynamics (4, 32). Fisher and Shannon information are related - Shannon information represents a coarse-graining (as below) of Fisher(35).

Here we investigate chemical reactions as both a physical and informational process. Consider an elemental reaction, A + B → C. As demonstrated in Figure 1, chemical reactions can be framed as a process governed by Gibbs free energy (36). For the reaction to proceed, the product(s) must have a lower free energy than the reactants with a resulting loss of heat. However, a thermodynamically favorable reaction must also proceed through an interaction channel that governs the *rate* of the reaction. Empirically, the rate of reaction is observed to be dependent on temperature as expressed in the Arrhenius equation:

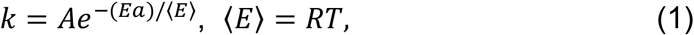

Where *k* is the rate, *A* is a “pre-exponential factor”, *E*_*a*_ is the “activation energy”, *R* is the universal gas constant, and *T* is absolute temperature (in K_o_). A critical property of the Arrhenius is the assumption that the activation energy (*E*_*a*_) is independent of the thermodynamic state of the system. As a result, an increase in system temperature increases the probability the energy of the reactants will exceed *E*_*a*_ so that the reaction is accelerated. Conventionally, enzymes are viewed as acting on the system by reducing *E*_*a*_.

The Arrhenius equation has evolved extensively since its conception with integration of Gibbs free energy, Boltzmann mechanics, and quantum mechanics in a variety of reformulations and re-interpretations. In collision theory, the probability of a reaction to be dependent on the energy with which the reactants collide. This takes on a statistical function because the actual reactant kinetic energy is determined by a Boltzmann distribution around some mean value at any given temperature (Figure 2). Thus, the probability an impact of two potential reactants will exceed *E*_*a*_ increases with the temperature.

Of interest here is the “pre-exponential factor” (*A*) which can be viewed as a “frequency factor” in collision theory (i.e., the frequency and orientation of molecular collisions) or as the “entropy of activation” in Transition State Theory. Here we restate this by viewing *A* as some probability that the interaction of the reactants will occur at some optimal quantum orientation *θ*that allows the reaction to occur.

Commonly, the actions of an enzyme are interpreted as a decrease in the activation energy (*E*_*a*_) (Figure 1). However, as noted above, the mechanism enzyme acceleration of a reaction typically requires “capture” of the reactants (i.e., reducing their velocity and kinetic energy) and placement in an orientation that optimizes overlap of their quantum wave functions and thus increase the probability of a reaction. Thus, an enzyme, by increasing the probability of an interaction between the substrate will result in a reaction, would appear to primarily alter the value of the pre-exponential factor (*A*) in the classical Arrhenius equation.

As we have noted previously, this probability function allows the reaction channel to be analyzed as an information-theoretic process subject to the principle of Maximum Fisher Information (MFI), which is a conservation law that requires Fisher information loss (as heat) in any physical process. This principle has been used to derive most of equations in physics including the Arrhenius equation(29, 32, 34, 37). Simply stated, the MFI principles requires the FI that enters a reaction channel to exceed the FI of the product(s). Here, the FI is broadly defined by the position and momentum space of the interacting reactants(22, 23) which also governs the probability of the reaction (Figure 3). A similar approach can be applied to the role of temperature on the rate of interactions when the probability of a reaction requires the kinetic energy of the reactants (now described by a Boltzmann distribution around some mean) exceeds the activation energy (Figure 2).

Thus, in the absence of a catalyst, A + B interactions represents a probability function defined by the Boltzmann distribution of the kinetic energies of each reactant – determined by the system temperature. As the temperature increases, the mean energy of the interactions of the reactants increases so that the corresponding probability of a reaction and FI also increases producing an acceleration of the reaction rate in the system.

Here we focus on the dynamics by which an enzyme increases the FI of the reactants without altering their kinetic energy. This can be analyzed mathematically as follows:

We use *p*(***x***) to quantify the FI in the interactions of the reactants. Mathematically *p*(***x***) is a probability law governing the properties of the A and B interactions ***x***, which are any consistently observable phenomenon. Here, *p*(***x***) includes the kinetic energy *E* of the two reactants (A and B), which is dependent on the system temperature. The *p*(***x***) also includes the physical orientation *θ*at which they interact. As a simplifying assumption, we view these as independent processes – that is, an enzyme captures each reactant molecule reducing its kinetic energy to 0 so that its role is entirely on optimize *θ*. In reality, this assumption may not be entirely correct – for example, the size and flexibility of enzymes may permit them to impact the substrate at a non 0 kinetic energy.

The relationship of the informational state of the enzymatic interactions with the kinetic energy of non-catalyzed reactions can be quantified (32, 33, 38) using the following equation that restates the chemical reaction as a communication channel with the coordinates of energy and a possible catalysts *F* which can alter either the energy (E) of their interactions or by optimizing their quantum orientation (*θ*) and an output molecule *C*.

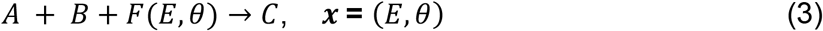

This channel communicates Fisher information about its reacting molecules *A, B* to C through the catalyzation *property F*. Specifically, the input information is carried in coordinates ***x* =** (*E, θ*), with *E* the *total* reactant energy of molecules *A, B*; and *θ*the angle of relative orientation between them which we derive from coarse graining the actual quantum interactions (see below).

Mathematically, the reaction channel has a solution *p*(***x***) = *p*(*E, θ*) in “energy-orientation” space describing the output for any given values for coordinates ***x*** ≡ *E, θ*.

### Coarse graining to link Fisher information in the reaction and Shannon information in the genome

The reaction channel in A + B → C in Eq. (3) involves dynamics at multiple spatial scales. At finer scales the probability of a reaction is governed by the quantum states of the electrons forming the bonds at these positions defined by the von Neumann entropy (13). At observable spatial scales, the reaction rate is governed by the Arrhenius equation and the kinetic energy of the molecular reactants and the frequency of their interactions. Bridging these scales requires “coarse graining.”

Coarse graining allows system structure that would, otherwise, be “missed” to become prominent. These have found practical applications in molecular dynamics and here we use this approach to identify the unobservable quantum angular *θ*dependence of our reaction. In coarse-graining, differential distances are replaced by finite displacements *d****x***→ Δx. A consequence of this approach is that the Fisher information is approximated by the Shannon information. Information form Eq. (1) is then used to describe the reaction dynamics by using the total energy *E* and angle *θ*between two chemical reactants as the two *x*_*n*_ coordinates.

### Fisher information and cellular thermodynamics

The form Eq. (1) for the Fisher Information holds most generally on a *continuous* space, i.e., of points ***x*** = *E, θspaced* by amounts *d****x*** → 0. However, in our *coarse-grained* space, with effective grain increment, or simply “size” *d****x*** → Δ***x*** *finite*, it is well-approximated by

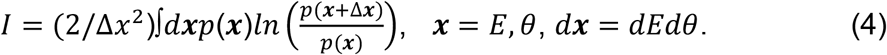

These grain *increments* are effective “smudges” of uncertainty that, in a quantum interpretation, represent actual quantized increments. Here, we view coarse graining as a spatial process – the information encoded in the genome and that result in molecular interactions is much larger than the quantum-level dynamics that govern the wave function interactions that govern the reaction. Note the Fisher information Eq. (1) has become the classical Kullback-Leibler (39) *cross entropy* between the occurrence law *p*(***x***) and its shifted version *p*(***x*** + Δ***x***). This is also equivalent to the Shannon information (4). In prior work, we have shown the key-lock spatial matching of an enzyme and substrate is quantified by Kullback-Leibler cross entropy (29). Optimal spatial matching produces a minimization of the K-L entropy.

The coarseness Δ*x* of the spacing Δx in K-L Eq. (4) renders it a quasi-*thermodynamic* quantity. That is, although p(***x***) is defined algebraically for a generally quantum effect as |a(***x***)exp(i*∅*(***x***)|^2^, with *∅*(***x***) the usual quantum phase, the information evaluated as Eq. (4) becomes manifestly ‘blind’ to these *phase values ∅*(***x***) (they drop out). However, as we will see, it is *not* blind to geometrical *orientation* angles θ or their increments Δθ. These may be considered as “residual” quantum phase effects. In other words, coarse graining allows us to transition from quantum interactions to macroscopic properties including the substrate molecular angle of interaction.

Thus, working with a mathematically smoothed (coarse-grained) channel loses some FI in pre-existing quantum phase information converting Fisher Information to Shannon entropy. However, coarse graining provides a natural route to derivation of (a) the Arrhenius exponential p(E) on total energy *E* in a two-reactant chemical system; and, independently, (b) a “Knudsen-type” complex exponential (cosine) form p(θ). Furthermore, the chemical system dynamics now be expressed in the metric (i.e., K-L entropy or Shannon information) that also describes genetic information.

### State of thermodynamic equilibrium

To review, coordinates ***x*** are now *E, θ*. These are the *total* energy levels *E* ≡ *E*_*A*_ + *E*_*B*_ of the two reactants, and the quantum angles *θ*of relative orientation between *A* and *B* at their interaction. We note there are inefficiencies inherent to both processes. For a reaction to proceed, the energy E must exceed the transition energy of the reaction, which is greater than the sum of the mean energy of each reactant. Similarly, the probability of reaction decreases as the angle of interaction diverges from some optimal value *θ*_*0*._

Here we assume that the enzyme binding to the substrate molecules effectively reduce their kinetic energy to 0 so that the reaction rate is dependent upon *p*(*E, θ*) where *E* and *θ*are independent variables.

Thus, for an enzyme, maximal reaction acceleration requires the minimum separation of reactant quantum states which we view as minimum divergence from some optimal value of θ. This can be restated as a *minimum loss* of information so that the system law *p*(***x***) obeys

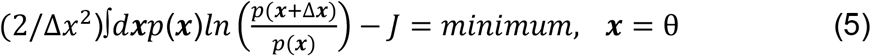

at all such reactions. As we saw, this is now a *thermodynamic* requirement on the K-L cross-entropy, which also represents the *Shannon information I*. Therefore, from the required of a minimum in Eq. (5) for its value from *J*, it also represents a condition of maximum Fisher information. The solution to (5) is found in Supplemental Material

## Acknowledgements

This work was supported by the Moffitt Physical Science Oncology Center NIH grant U54 CA143970. The author thanks Chris Whelan and Joel Brown for the valuable editorial advice.

## Author contribution

This is a single authored manuscript.

## Data Availability

All data generated or analyzed during this study are included in this published article [and its supplementary information files].

